# Unsupervised multiple kernel learning for heterogeneous data integration

**DOI:** 10.1101/139287

**Authors:** Jérôme Mariette, Nathalie Villa-Vialaneix

## Abstract

Recent high-throughput sequencing advances have expanded the breadth of available omics datasets and the integrated analysis of multiple datasets obtained on the same samples has allowed to gain important insights in a wide range of applications. However, the integration of various sources of information remains a challenge for systems biology since produced datasets are often of heterogeneous types, with the need of developing generic methods to take their different specificities into account.

We propose a multiple kernel framework that allows to integrate multiple datasets of various types into a single exploratory analysis. Several solutions are provided to learn either a consensus meta-kernel or a meta-kernel that preserves the original topology of the datasets. We applied our framework to analyse two public multi-omics datasets. First, the multiple metagenomic datasets, collected during the *TARA* Oceans expedition, was explored to demonstrate that our method is able to retrieve previous findings in a single KPCA as well as to provide a new image of the sample structures when a larger number of datasets are included in the analysis. To perform this analysis, a generic procedure is also proposed to improve the interpretability of the kernel PCA in regards with the original data. Second, the multi-omics breast cancer datasets, provided by The Cancer Genome Atlas, is analysed using a kernel Self-Organizing Maps with both single and multi-omics strategies. The comparison of this two approaches demonstrates the benefit of our integration method to improve the representation of the studied biological system.

Proposed methods are available in the R package **mixKernel**, released on CRAN. It is fully compatible with the **mixOmics** package and a tutorial describing the approach can be found on **mixOmics** web site http://mixomics.org/mixkernel/.

## 1 Introduction

Recent high-throughput sequencing advances have expanded the breadth of available omics datasets from genomics to transcriptomics, proteomics and methylomics. The integrated analysis of multiple datasets obtained on the same samples has allowed to gain important insights in a wide range of applications from microbial communities profiling [14] to the characterization of molecular signatures of human breast tumours [35]. However, multiple omics integration analyses remain a challenging task, due to the complexity of biological systems, heterogeneous types (continuous data, counts, factors, networks…) between omics and additional information related to them and the high-dimensionality of the data.

In the literature, several strategies have been proposed to analyse multi-omics datasets. Multivariate approaches is a widely used framework to deal with such problems and several such methods (including PLS and CCA) are provided in the R package **mixOmics** [19]. Similarly, the multiple co-inertia analysis [8], use the variability both within and between variables to extract the linear relationships that best explain the correlated structure across datasets. However, these approaches are restricted to the analysis of continuous variables and thus are not generic in term of data types used as inputs. Some works use a case-by-case approach to integrate non numeric information into the analysis: [42] propose a joint non-negative matrix factorization framework to integrate expression profiles to interaction networks by adding network-regularized constraints with the help of graph adjacency matrices and [12,26] propose extensions of the widely used PCoA approach to integrate information about phylogeny and environmental variables. Finally, some authors propose to use a transformation of all the input datasets into a unified representation before performing an integrated analysis: [16] transforms each data types into graphs, which can be merged before being analysed by standard graph measures and graph algorithms. However, graph based representation are a very constraining and rough way to represent a complex and large dataset.

In the present work, we take advantage of the kernel framework to propose a generic approach that can incorporate heterogeneous data types as well as external information in a generic and very flexible way. More precisely, any dataset is viewed through a kernel, that provides pairwise information between samples. Kernels are a widely used and flexible method to deal with complex data of various types: they can be obtained from *β*-diversity measures [5,23] to explore microbiome datasets. They can also account for datasets obtained as read counts by the discrete Poisson kernel [7] and are also commonly adopted to quantifies genetic similarities by the state kernel [17,39]. Our contribution is to propose three alternative approaches able to combine several kernels into one meta-kernel in an unsupervised framework. If multiple kernel approaches are widely developed for supervised analyses, unsupervised approaches are less easy to handle, because no clear *a priori* objective is available. However, they are required to use kernel in exploratory analyses that are the first step to any data analysis.

To evaluate the benefits of the proposed approach, two datasets have been analysed. The first one is the multiple metagenomic dataset collected during the *TARA* Oceans expedition [3,15] and the second one is based on a multi-omic dataset on breast cancer [35]. A method to improve the interpretability of kernel based exploratory approaches is also presented and results show that not only our approach allows to retrieve the main conclusions stated in the different papers in a single and fast analysis, but that it can also provide new insights on the data and the typology of the samples by integrating a larger number of information.

## 2 Methods

### 2.1 Unsupervised multiple kernel learning

#### 2.1.1 Kernels and notations

For a given set of observations (*x*_*i*_)_*i*=1_,…_*N*_, taking values in an arbitrary space 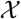, we call “kernel” a function 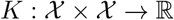 that provides pairwise similarities between the observations: *K*_*ij*_:= *K*(*x*_*i*_,*x*_*j*_). Moreover, this function is assumed to be symmetric (*K*_*ij*_ = *K*_*ji*_) and positive

(∀*n* ∈ ℕ, ∀(*α*_*i*_)_*i*=1,…,*n*_ ⊂ ℝ, 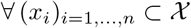, 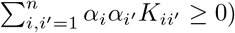. According to [1], this ensures that *K* is the dot product in a uniquely defined Hilbert space (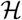, 〈.,.〉) of the images of (*x*_*i*_)_*i*_ by a uniquely defined feature map *ϕ*: 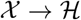: *K*_*ij*_ = 〈*ϕ*(*x*_*i*_),*ϕ*(*x*_*j*_)〉. In the sequel, the notation *K* will be used to denote either the kernel itself or the evaluation matrix (*K*_*ij*_)_*i*,*j*=1,…,*N*_ depending on the context.

This setting allows us to deal with multiple source datasets in a uniform way, provided that a relevant kernel can be calculated from each dataset (examples are given in Section 3.1 for standard numeric datasets, phylogenetic tree, …). Suppose now that *M* datasets 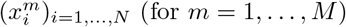 are given instead of just one, all obtained on the same samples *i* = 1,…, *N*. *M* different kernels (*K*^*m*^)_*m*=1,…,*M*_ provide different views of the datasets, each related to a specific aspect.

Multiple kernel learning (MKL) refers to the process of linearly combining the *M* given kernels into a single kernel *K*\*:

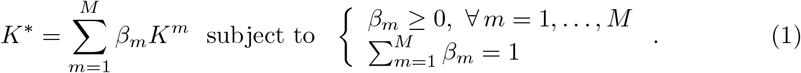

By definition, the kernel *K*\* is also symmetric and positive and thus induces a feature space and a feature map (denoted by *ϕ*\* in the sequel). This kernel can thus be used in subsequent analyses (SVM, KPCA, KSOM, …) as it is supposed to provide an integrated summary of the samples.

A simple choice for the coefficients *β*_*m*_ is to set them all equal to 1/*M*. However, this choice treats all the kernels similarly and does not take into account the fact that some of the kernels can be redundant or, on the contrary, atypical. Sounder choices aim at solving an optimization problem so as to better integrate all informations. In a supervised framework, this mainly consists in choosing weights that minimize the prediction error [13]. For clustering, a similar strategy is used in [40], optimizing the margin between the different clusters. However, for other unsupervised analyses (such as exploratory analysis, KPCA for instance), such criteria do not exist and other strategies have to be used to choose relevant weights.

As explained in [41], propositions for unsupervised multiple kernel learning (UMKL) are less numerous than the ones available for the supervised framework. Most solutions (see, *e.g.*, [22,41]) seek at providing a kernel that minimizes the distortion between all training data and/or that minimizes the approximation of the original data in the kernel embedding. However, this requires that the datasets 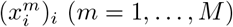 are standard numerical datasets: the distortion between data and the approximation of the original data are then directly computed in the input space (which is ℝ^*d*^) using the standard Euclidean distance as a reference. Such a method is not applicable when the input dataset is not numerical (*i.e.*, is a phylogenetic tree for instance) or when the different datasets 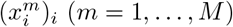 do not take value in a common space.

In the sequel, we propose two solutions that overcome this problem: the first one seeks at proposing a consensual kernel, which is the best consensus of all kernels. The second one uses a different point of view and, similarly to what is suggested in [41], computes a kernel that minimizes the distortion between all training data. However, this distortion is obtained directly from the *M* kernels, and not from an Euclidean input space. Moreover, it is used to provide a kernel representation that preserve the original data topology. Two variants are described: a sparse variant, which also selects the most relevant kernels, and a non sparse variant, when the user does not want to make a selection among the *M* kernels.

#### 2.1.2 A consensus multiple kernel

Our first proposal, denoted by STATIS-UMKL, relies on ideas similar to STATIS [18,20]. STATIS is an exploratory method designed to integrate multi-block datasets when the blocks are measured on the same samples. STATIS finds a consensus matrix, which is obtained as the matrix that has the highest average similarity with the relative positions of the observations as provided by the different blocks. We propose to use a similar idea to learn a consensus kernel.

More precisely, a measure of similarity between kernels can be obtained by computing their cosines1 according to the Frobenius dot product: ∀*m*, *m*′ = 1, …, *M*,

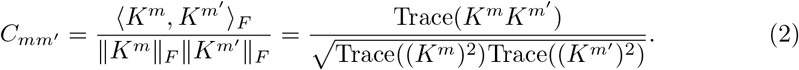

*C*_*mm′*_ can be viewed as an extension of the RV-coefficient [30] to the kernel framework, where the RV-coefficient is computed between 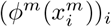 and 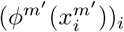 (where *ϕ*^*m*^ is the feature map associated to *K*^*m*^).

The similarity matrix **C** = (*C*_*mm′*_)_*m,m′*=1,…,*M*_ provides information about the resemblance between the different kernels and can be used as such to understand how they complement each other or if some of them provide an atypical information. It also gives a way to obtain a summary of the different kernels by choosing a kernel *K*\* which maximizes the average similarity with all the other kernels:

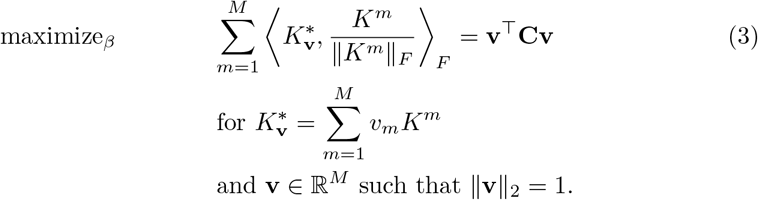

The solution of the optimization problem of Equation (3) is given by the eigen-decomposition of **C**. More precisely, if **v** = (*v*_*m*_)_*m*=1,…,*M*_ is the first eigenvector (with norm 1) of this decomposition, then its entries are all positive (because the matrices *K*^*m*^ are positive) and are the solution of the maximization of **v**^Τ^**Cv**. Setting 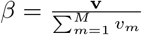 thus provides a solution satisfying the constrains of Equation (1) and corresponding to a consensual summary of the *M* kernels.

Note that this method is equivalent to performing multiple CCA between the multiple feature spaces, as suggested in [38] in a supervised framework, or in [28] for multiple kernel PCA. However, only the first axis of the CCA is kept and a *L*^2^-norm constrain is used to allow the solution to be obtained by a simple eigen-decomposition. This solution is better adapted to the case where the number of kernels is small.

#### 2.1.3 A sparse kernel preserving the original topology of the data

Because it focuses on consensual information, the previous proposal tends to give more weights to kernels that are redundant in the ensemble of kernels and to discard the information given by kernels that provide complementary informations. However, it can also be desirable to obtain a solution which weights the different images of the dataset provided by the different kernels more evenly. A second solution is thus proposed, which seeks at preserving the original topology of the data. This method is denoted by sparse-UMKL in the sequel.

More precisely, weights are optimized such that the local geometry of the data in the feature space is the most similar to that of the original data. Since the input datasets are not Euclidean and do not take values in a common input space, the local geometry of the original data cannot be measured directly as in [41]. It is thus approximated using only the information given by the *M* kernels. To do so, a graph, the *k*-nearest neighbour graph (for a given *k* ∈ ℕ*), 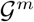, associated with each kernel *K*^*m*^ is built. Then, a (*N* × *N*)-matrix **W**, representing the original topology of the dataset is defined such that *W*_*ij*_ is the number of times the pair (*i,j*) is in the edge list of 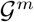 over *m* = 1,…,*m* (*i.e.*, the number of times, over *m* = 1,…,*M*, that 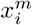 is one of the *k* nearest neighbours of 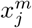 or 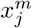 is one of the *k* nearest neighbours of 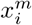).

The solution is thus obtained for weights that ensure that *ϕ*\*(*x*_*i*_) and *ϕ*\*(*x*_*j*_) are “similar” (in the feature space) when *W*_*ij*_ is large. To do so, similarly as [22], we propose to focus on some particular features of *ϕ*\*(*x*_*i*_) which are relevant to our problem and correspond to their similarity (in the feature space) with all the other *ϕ*\*(*x*_*j*_). More precisely for a given *β* ∈ ℝ^*M*^, we introduce the *N*-dimensional vector 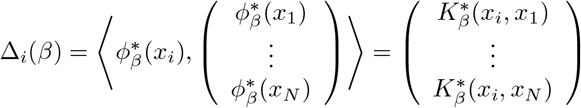. But, contrary to [22], we do not rely on a distance in the original space to measure topology preservation but we directly use the information provided by the different kernels through **W**. The following The optimization problem is thus solved:

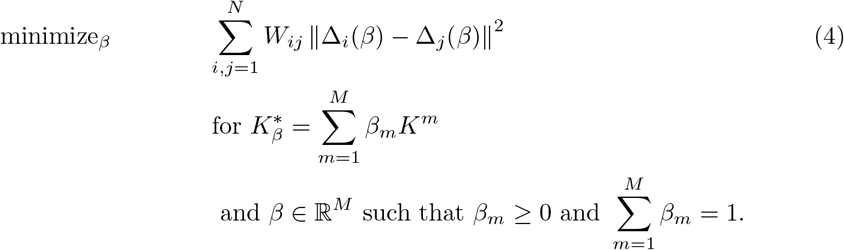

The optimization problem of Equation (4) expands as

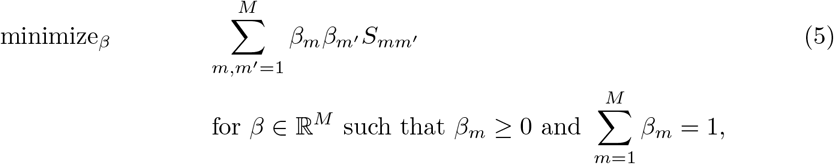

for 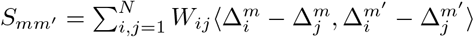 and 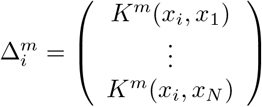. The matrix **S** = (*S*_*m,m′*=1,…,*M*_ is positive and the problem is thus a standard Quadratic Programming (QP) problem with linear constrains, which can be solved by using the R package **quadprog**. Since the constrain 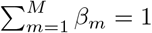 is an *L*^1^ constrain in a QP problem, the produced solution will be sparse: a kernel selection is performed because only some of the obtained (*β*_*m*_)_*m*_ are non zero. While desirable when the number of kernels is large, this property can be a drawback when the number of kernels is small and that using all kernels in the integrated exploratory analysis is expected. To address this issue, a modification of Equation (5) is proposed in the next section.

#### 2.1.4 A full kernel preserving the original topology of the data

To get rid of the sparse property of the solution of Equation (5), an *L*^2^ constrain can be used to replace the *L*^1^ constrain, similarly to Equation (3):

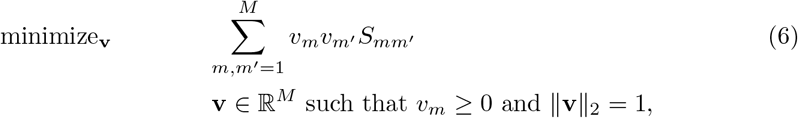

and to finally set 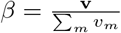. This problem is a Quadratically Constrained Quadratic Program (QCQP), which is known to be hard to solve. For a similar problem, [22] propose to relax the problem into a semidefinite programming optimization problem. However, a simpler solution is provided by using ADMM (Alterning Direction Method of Multipliers; [4]). More precisely, the optimization problem of Equation (6) is re-written as

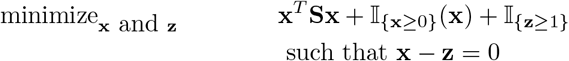

and is solved with the method of multipliers. Final weights are then obtained by re-scaling the solution 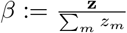. The method is denoted by full-UMKL in the sequel.

### 2.2 Kernel PCA (KPCA) and enhanced interpretability

The combined kernel can be used in subsequent exploratory analyses to provide an overview of the relations between samples through the different integrated datasets. Any method based only on dot product and norm computations can have a kernel version and this includes a variety of standard methods, such as PCA (KPCA, see below), clustering (kernel *k*-means, [32]) or more sophisticated approaches that combine clustering and visualization like self-organizing maps (kernel SOM, [24]). In this section, we focus on the description of KPCA because it is close to the standard approaches that are frequently used in metagenomics (PCoA) and is thus a good baseline analysis for investigating the advantages of our proposals. Moreover, we have used KPCA to propose an approach that is useful to improve the interpretability of the results. Section 4.2 illustrates that our method is not restricted to this specific analysis and is straightforwardly extensible to other exploratory tools.

#### 2.2.1 Short description of KPCA

KPCA, introduced in [31], is a PCA analysis performed in the feature space induced by the kernel *K*\*. It is equivalent to standard MDS (*i.e.*, metric MDS or PCoA; [36]) for Euclidean dissimilarities. Without loss of generality, the kernel *K*\* is supposed centered 2. KPCA simply consists in an eigen-decomposition of *K*\*: if (*α*_*k*_)_*k*=1,…,*N*_ ∈ ℝ^*N*^ and (λ_*k*_)_*k*=1,…,*N*_ respectively denote the eigenvectors and corresponding eigenvalues (ranked in decreasing order) then the PC axes are, for *k* = 1, …, *N*, 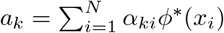, where *α*_*k*_ = (*α*_*ki*_)_*i*=1,…,*N*_. **a**_*k*_ = (a_*ki*_)_*i*=1,…,*N*_ are orthonormal in the feature space induced by the kernel: ∀ *k*, *k′*, 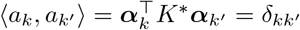 with 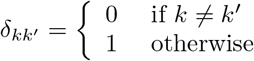. Finally, the coordinates of the projections of the images of the original data, (*ϕ*\*(*x*_*i*_))_*i*_, onto the PC axes are given by: 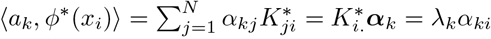, where 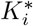 is the *i*-th row of the kernel *K*\*.

These coordinates are useful to represent the samples in a small dimensional space and to better understand their relations. However, contrary to standard PCA, KPCA does not come with a variable representation, since the samples are described by their relations (via the kernel) and not by standard numeric descriptors. PC axes are defined by their similarity to all samples and are thus hard to interpret.

#### 2.2.2 Interpretation

There are few attempts, in the literature, to help understand the relations of KPCA with the original measures. When the input datasets take values in ℝ^*d*^, [29] propose to add a representation of the variables to the plot, visualizing their influence over the results from derivative computations. However, this approach would make little sense for datasets like ours, *i.e.*, described by discrete counts.

We propose a generic approach that assesses the influence of variables and is based on random permutations. More precisely, for a given measure *j*, that is used to compute the kernel *K*^*m*^, the values observed on this measure are randomly permuted between all samples and the kernel is re-computed: 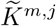. For species abundance datasets, the permutation can be performed at different phylogeny levels, depending on the user interest. Then, using the weights found with the original (non permuted) kernels, a new meta-kernel is obtained 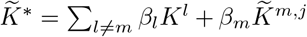. The influence of the measure *j* on a given PC subspace is then assessed by computing the Crone-Crosby distance [9] at the axis level: ∀ *k* = 1, …, *N*, 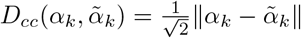, where *α*_*k*_ and 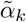 respectively denote the eigenvectors of the eigen-decomposition of *K*\* and 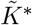. 3.

Finally, the KPCA interpretation is done similarly as for a standard PCA: the interpretation of the axes (*a*_*k*_)_*k*=1,…,*N*_ is done with respect to the observations (*x*_*i*_)_*i*=1,…,*N*_ which contribute the most to their definition, when important variables are the ones leading to the largest Crone-Crosby distances.

Methods presented in the paper are available in the R package **mixKernel**, released on CRAN. Further details about implemented functions are provided in Supplementary Section S1.

## 3 Case studies

### 3.1 *TARA* Oceans

The *TARA* Oceans expedition [3, 15] facilitated the study of plankton communities by providing oceans metagenomic data combined with environmental measures to the scientific community. During the expedition, 579 samples were collected for morphological, genetic and environmental analyses, from 75 stations in epipelagic and mesopelagic waters across eight oceanic provinces. The *TARA* Oceans consortium partners analysed prokaryotic [34], viral [6] and eukaryotic-enriched [11] size fractions and provided an open access to the raw datasets and processed materials. So far, all articles related to *TARA* Oceans that aim at integrating prokaryotic, eukaryotic and viral communities, took advantage of the datasets only by using co-occurrence associations [14, 21, 37]. The integration analysis of the whole material aims at providing a more complete overview of the relations between all collected informations.

48 selected samples were collected in height different oceans or seas: Indian Ocean (IO), Mediterranean Sea (MS), North Atlantic Ocean (NAO), North Pacific Ocean (NPO), Red Sea (RS), South Atlantic Ocean (SAO), South Pacific Ocean (SPO) and South Ocean (SO). Using these samples, 8 (dis)similarities were computed using public preprocessed datasets, which are all available from the *TARA* Oceans consortium partners websites. These dissimilarities provide information about environmental variables, phylogenetic similarities, prokaryotic functional processes, different aspects of the eukaryotic dissimilarities and virus composition. Selected datasets as well as chosen kernels are fully described in Supplementary Section S2. The meta-kernel was analysed using a KPCA and the most important variables were assessed as described in Section 2.2.2.

### 3.2 TCGA

[35] (TCGA) provides multi-omics datasets from different tumour types, such as colon, lung and breast cancer. In this work, we consider normalized and pre-filtered breast cancer datasets available from the **mixOmics** website4. Using the 989 available samples, three kernels were computed. The **TCGA.mRNA** kernel provides a gene expression dissimilarity measure computed on the expression of 2,000 mRNAs, the **TCGA.miRNA** describes the expression of 184 miRNAs and the methylation aspect is assessed by the **TCGA.CpG** kernel, computed on 2,000 CpG probes. These three kernels were obtained using the Gaussian kernel, 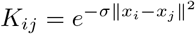 with σ equal to the median of 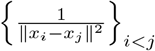. The combined kernel was used in kernel self-organizing map (KSOM) that has been implemented with the R package **SOMbrero** [2]. KSOM is an exploratory tool that combines clustering and visualization by mapping all samples onto a 2-dimensional (generally squared) grid made of a finite number of units (also called clusters). It has been shown relevant, *e.g.* to provide a relevant taxonomy of Amazonian butterflies from DNA barcoding in [25].

The results of our analysis with the combined kernel were compared to the results obtained with a simple analysis that uses only one of the kernel. The comparison was performed using a quality measure specific to SOM, the *topographic error* (TE), which is the ratio of the second best matching unit that falls in the direct neighbor, on the grid, of the chosen unit over all samples [27]. In addition, breast cancer subtypes, *i.e., Basal, Her2, LumA* or *LumB* are provided for every sample and were used as an *a priori* class to compute clustering quality measures (they were thus excluded from the exploratory analysis). More precisely, (i) the average cluster purity, *i.e.*, the mean over all clusters on the grid of the frequency of the majority vote cancer subtype and (ii) the normalized mutual information (NMI) [10] between cancer subtype and cluster, which is a value comprised between 0 and 1 (1 indicating a perfect matching between the two classifications).

## 4 Results and discussion

Sections 4.1 and 4.2 provide and discuss results of exploratory analyses performed from the two sets of datasets described in the previous section. More precisely, Section 4.1 explores the datasets studied in [34], [6] and [11] with KPCA. This illustrates how a multiple metagenomic dataset can be combined with external information to provide an overview of the structure of the different samples. In addition, Section 4.2 shows that our approach is not restricted nor to metagenomic neither to KPCA by studying the multi-omic dataset related to breast cancer with KSOM.

All analyses presented in this section use the full-UMKL strategy. However, for both datasets, a study of the correlation between kernels in the line of the STATIS-UMKL approach is provided in Supplementary Section S4 and shows how this approach helps understand the relations between the multiple datasets. Moreover, a comparison between the different multiple kernel strategies is discussed in Supplementary Section S5 and justifies the choice of full-UMKL for our problems. All combined kernels have been implemented with our package **mixKernel**, as well as all KPCA results.

### 4.1 Exploring *TARA* oceans datasets with a single KPCA

In a preliminary study (fully reported in Supplementary Section S3), an exploratory analysis was performed using a KPCA with only the three *TARA* Oceans datasets studied in [34] and the full-UMKL strategy. The results show that the main variability between the different samples is explained similarly as in [34]: the most important variables returned by our method are those discussed in this article to state the main conclusions.

A further step is then taken by integrating all *TARA* Oceans datasets described in Supplementary Section S2. Supplementary Section S4.1 shows that **pro.phylo** and **euk.pina** are the most correlated kernels to environmental and physical variables, unlike large organism size fractions, *i.e.*, **euk.meso** which are strongly geographically structured. Figure 1 (left) displays the projection of the samples on the first two axes of the KPCA. Figure 1 (right) and Supplementary Figure S16 provide the 5 most important variables for each datasets, respectively for the first and the second axes of the KPCA. To obtain these figures, abundance values were permuted at 56 prokaryotic phylum levels for the **pro.phylo** kernel, at 13 eukaryotic phylum levels for **euk.pina**, **euk.nano**, **euk.micro** and **euk.meso** and at 36 virus family levels for the **vir.VCs** kernel. Variables used for **phychem** and **pro.NOGs** were the same than the one used in the restricted analysis. Additionally, the explained variance supported by the first 15 axes is provided in Supplementary Figure S17. Using an R implementation of the methods on a 1 CPU computer with 16GB memory, the computational cost to combine the three kernels is only ~3 seconds. Permutations to assess the eight kernels important variables are computationally much more demanding if performed at a fine level as we did. In our case, they took ~13 minutes.

**Figure 1.**
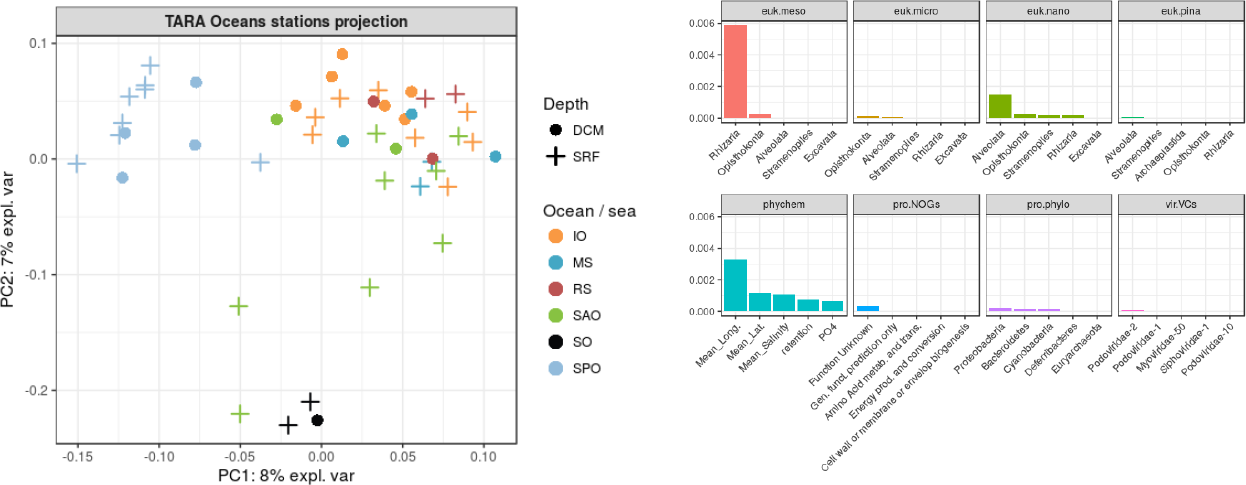
Left: Projection of the observations on the first two KPCA axes. Colors represent the oceanic regions and shapes the depth layers. Right: The 5 most important variables for each of the eight datasets, ranked by decreasing Crone-Crosby distance.

Contrary to the restricted analysis, Figure 1 does not highlight any particular pattern in terms of depth layers but it does in terms of geography. SO samples are gathered in the bottom-center of the KPCA projection and SPO samples are gathered on the top-left side. Figure 1 shows that the most important variables come from the **phychem** kernel (especially the longitude) and from kernels representing the eukaryotic plankton. More specifically, large size organisms are the most important: *rhizaria* phylum for **euk.meso** and *alveolata* phylum for **euk.nano**. The abundance of *rhizaria* organisms also ranks first between important variables of the second KPCA axis, followed by the *opisthokonta* phylum for **euk.nano**. The display of these variables on the KPCA projection reveals a gradient on the first axis for both the *alveolata* phylum abundance (Supplementary Figure S18) and the longitude (Supplementary Figure S19) and on the second axis for *rhizaria* (Supplementary Figure S20) and *opisthokonta* (Supplementary Figure S21) abundances. This indicates that SO and SPO epipelagic waters mainly differ in terms of *Rhizarians* abundances and both of them differ from the other studied waters in terms of *alveolata* abundances.

The integration of *TARA* Oceans datasets shows that the variability between epipelagic samples is mostly driven by geography rather than environmental factors and that this result is mainly explained by the strong geographical structure of large eukaryotic communities. Studied samples were all collected from epipelagic layers, where water temperature does not vary much, which explains the poor influence of the prokaryotic dataset in this analysis.

### 4.2 Clustering breast cancer multi-omics datasets

KSOM was used to obtain a map from the three datasets presented in Section 3.2: mRNAs, miRNAs and methylation datasets. The results were compared to a single-omic analysis with the same method (KSOM). KSOM maps were trained with a 5 × 5 grid, 5,000 iterations in a stochastic learning framework and a Gaussian neighbourhood controlled with the Euclidean distance between units on the grid. The computational cost to combine the kernels was ~20 minutes and ~12 seconds were required to generate one map.

Table 1 reports KSOM performances obtained over 100 maps (mean and standard deviation) for the three kernels: **TCGA.mRNA**, **TCGA.miRNA** and **TCGA.CpG** and the meta-kernel, denoted **TCGA.all**. Results are reported in terms of average cluster purity and NMI, with respect to the cancer subtypes. All TE were found to be equal to 0. This indicates a good organization of the results on the grid, with respect to the topology of the original dataset as represented in the input kernel. Finally the map with the best NMI obtained for the meta-kernel is given in Figure 2.

**Table 1.** Performance results (average cluster purity and NMI with respect to the cancer subtype) of KSOM (average over 100 maps and standard deviation between parenthesis) for the three single-omics datasets: **TCGA.mRNA**, **TCGA.miRNA** and **TCGA.CpG** and the meta-kernel, **TCGA.all**.

|  | TCGA.mRNA | TCGA.miRNA | TCGA.CpG | TCGA.all |
| --- | --- | --- | --- | --- |
| purity | 0.67 (0.02) | 0.62 (0.02) | 0.66 (0.01) | **0.70** (0.02) |
| NMI | 0.31 (0.02) | 0.17 (0.01) | 0.18 (0.01) | **0.34** (0.01) |

**Figure 2.**
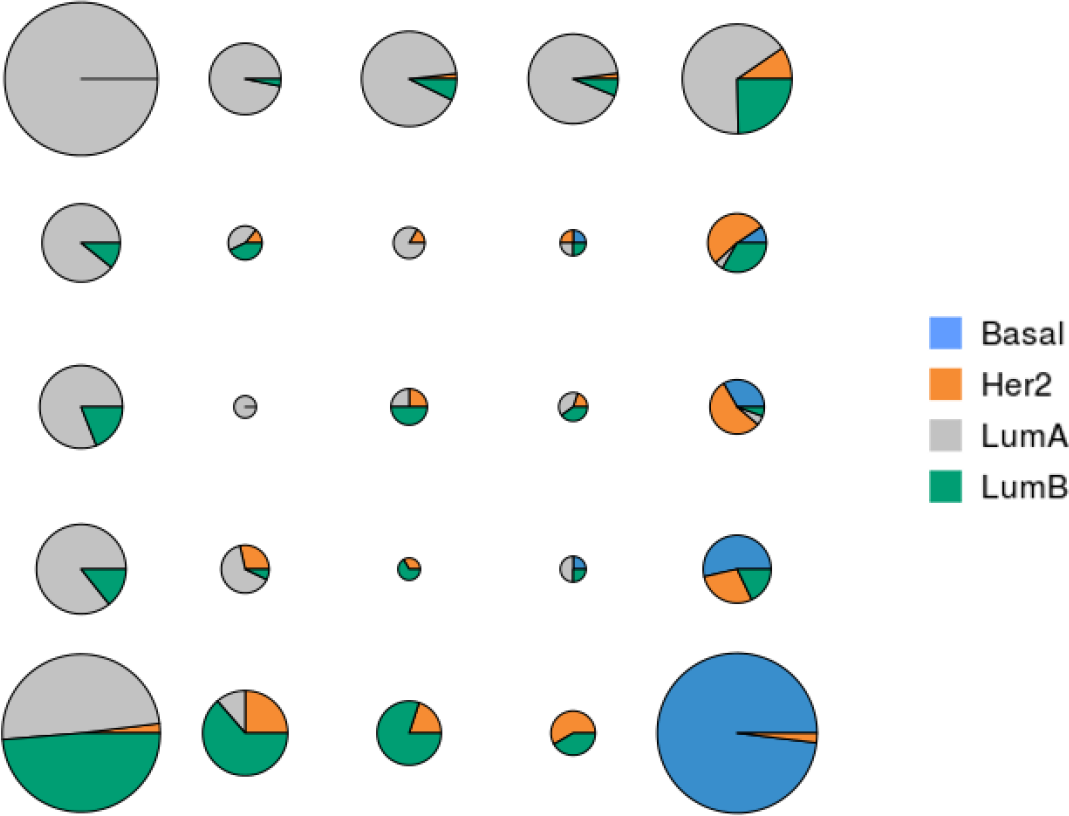
For each unit of the map, distribution of breast cancer subtypes. Colors represent the different breast cancer subtypes (*Basal*, *Her2*, *LumA* or *LumB*) and the area of the pie charts is proportional to the number of samples classified in the corresponding unit.

For all quality criteria, the integrated analysis gives better results (with respect to cancer subtype) than single-omics analyses (all differences are significant according to a student test, risk 1%). This can be explained by the fact that the information provided especially by mRNA and CpG are complementary, as described in the analysis of correlations between kernels in Supplementary Section S4.2. In addition, Figure 2 shows that the clustering produced by the KSOM is relevant to discriminate between the different breast cancer subtypes and to identify their relations (*e.g.*, subtypes *LumA* and *Basal* are closer to subtypes *LumB* and *Her2* than they are from each other). The organization of cancer subtypes on the map is in accordance with what is reported in [33] (from cDNA microarray). However, it has been obtained with additional datasets and thus provides a finer typology of samples. It shows that some *LumA* samples are mixed with *LumB* samples (cluster at the bottom left of the map) and that samples classified in the middle of the map probably have an ambiguous type. It also gives clue to define which samples are typical from a given cancer subtype. In addition, Supplementary Section S6 shows the results obtained by KPCA that are consistent with those of KSOM. It also provides a list of features (mRNA, miRNA and CpG probes) that are potentially interesting to discriminate between breast cancer subtypes.

## 5 Conclusion

The contributions of the present manuscript to the analysis of multi-omics datasets are twofolds: firstly, we have proposed three unsupervised kernel learning approaches to integrate multiple datasets from different types, which either allow to learn a consensual meta-kernel or a meta-kernel preserving the original topology of the data. Secondly, we have improved the interpretability of the KPCA by assessing the influence of input variables in a generic way.

The experiments performed on *TARA* Oceans and breast cancer datasets showed that presented methods allow to give a fast and accurate insight over the different datasets within a single analysis and is able to bring new insights as compared to separated single-dataset analyses. Future work include the addition of more kernels and post-processing methods for the analysis into our package **mixKernel**.

## 6 Acknowledgments

The authors are grateful to Rémi Flamary (University of Nice, France) for helpful suggestions on optimization and to the **mixOmics** team, especially to Kim-Anh Le Cao and Florian Rohart, for helping to integrate our method in the **mixOmics** package. We also thank the three anonymous reviewers for valuable comments and suggestions which helped to improve the quality of the paper.

## Footnotes

1 Cosines are usually preferred over the Frobenius dot product itself because they allow to re-scale the different matrices at a comparable scale. It is equivalent to using the kernel 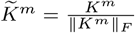 instead of *K*^*m*^.

2 if *K*\* is not centered, it can be made so by computing 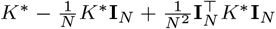, with **I**_*N*_ a vector with *N* entries equal to 1

3 Note that a similar distance can be computed at the entire projection space level but, since axes are naturally ordered in PCA, we chose to restrict to axis-specific importance measures.

4 http://mixomics.org/tcga-example/

